# Investigations into cyanobacteria, plant, and insect protein extracts as serum-replacement supplements for the expansion of cells for cultivated meat

**DOI:** 10.64898/2026.01.28.702276

**Authors:** William Gordon-Petrovskii, Georgie Hurst, Zara Dodhia, Paul Cameron, Michael Sulu, Gary J. Lye, Mariana Petronela Hanga

**Affiliations:** Department of Biochemical Engineering, University College London, London, WC1E 6BT, UK; Natural Resources Institute, University of Greenwich, Kent, UK; Manufacturing Futures Lab, University College London, London, E20 2AE, UK

**Keywords:** cultivated meat, serum-free media, food grade high protein extracts, spirulina, faba bean, mealworm

## Abstract

Cultivated meat has undeniable potential to address some of the current detrimental impacts of animal farming, while addressing food security worldwide. However, one of the main challenges in cultivated meat production is manufacturing cost. The main contributor to cost is the culture media which comprises expensive components such as growth factors and animal-derived proteins. This study investigated alternative, food grade, high protein extracts as serum replacements in serum-free media formulations. The extracts were chosen to represent various sustainable sources of proteins: marine (spirulina *e.g.* cyanobacterium), plant (faba bean) and insect (mealworm flour). Different processing methods and different solvents were investigated for production of cell culture-compatible extracts which were then tested with mouse myoblasts (C2C12) and primary porcine myosatellites (pMyoSCs). A serum-free medium formulation containing 2.6% v/v spirulina extract was found to support long term growth of C2C12 cells for ∼10 population doublings compared to only ∼2 in the control. The processing steps were optimized, showing that a glycerine solution was best for free amino acid and protein yield (4950 µM total free amino acids, 11.45 mg/mL protein concentration). This solution had a positive effect on C2C12 cells, increasing their growth by up to 20% when added to the B8 medium. However, this benefit did not translate to pMyoSCs, which showed no significant growth increases in short-term screening. This work demonstrates a method for converting food grade protein powders into effective culture media supplements and highlights the potential of spirulina-based extracts for the use in cultivated meat.

Graphical abstract
Created in BioRender. Gordon-Petrovskii, W. (2025) https://BioRender.com/by7khs1

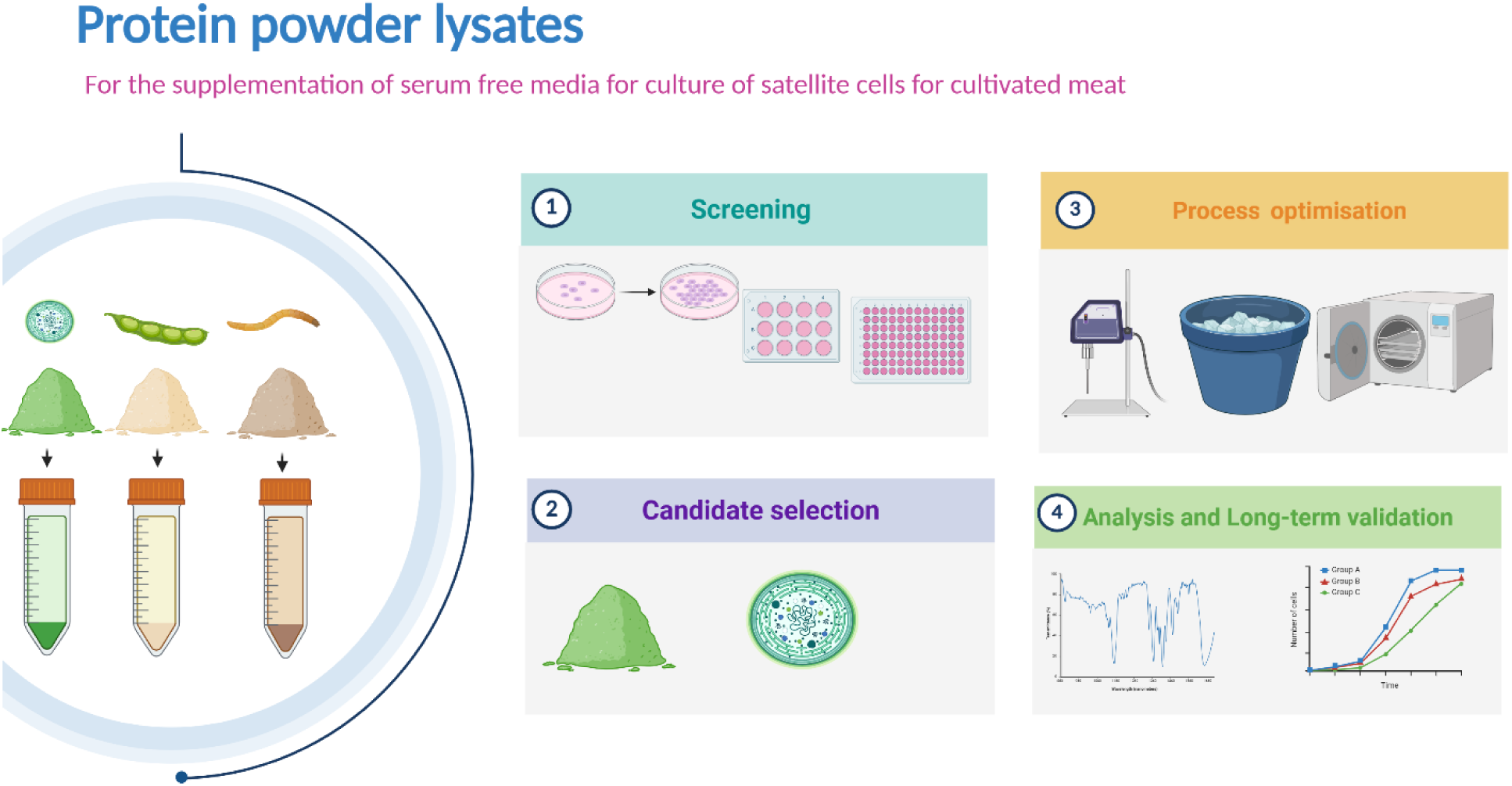

## Introduction

The demand for meat to feed a growing population are increasing (Beck, Haerlin, & Richter, 2016) leading to a need for sustainable solutions for meat production, while also addressing the negative impacts of current farming practices. Cultivated meat embraces the principles of animal cell culture and biotechnology to produce meat analogues without the slaughter of animals through the culture of cells *in vitro,* thus providing a more sustainable and ethically preferable alternative to conventional meat (Datar & Betti, 2010; Sinke, Swartz, Sanctorum, van der Giesen, & Odegard, 2023).

One key challenge for the development of cultivated meat technology is the transition away from the reliance on foetal bovine serum (FBS) in culture media. FBS is a commonly used media supplement which mimics the *in vivo* tissue microenvironment, by providing growth and attachment factors, fatty acids, hormones, antioxidants and other important compounds for cell maintenance and growth (D. Y. Lee et al., 2022). However, as it is derived from foetal cows whose mothers have been slaughtered whilst pregnant, there are therefore several ethical, environmental, and safety disadvantages associated with the use of FBS for cultivated meat production (Jochems, van der Valk, Stafleu, & Baumans, 2002). The food industry also faces significant cost limitations compared with the biopharmaceutical sector. Consequently, the cost of FBS is highly prohibitive. This issue is further compounded by its inconsistent supply which would be compounded if cultivated meat used FBS at large volumes due to its method of derivation (Dessels, Potgieter, & Pepper, 2016; Gstraunthaler, Lindl, & van der Valk, 2013). Therefore, the replacement of FBS with a preferably non-animal-derived alternative represents a pivotal step towards making cultivated meat a more viable, eco-conscious, and ethical choice for consumers.

Typical approaches for replacing FBS involve using defined components such as proteins like albumin, growth factors, minerals like selenium, insulin etc. Tissue culture research which replaces FBS typically uses pharmaceutical-grade ingredients for this application, however with a food product like cultivated meat, pharmaceutical-grade ingredients are too expensive, representing 95% of the manufacturing costs in the cultivated meat production (Specht, 2020). By utilising food-grade ingredients instead, there is potential to reduce the cost of serum-free media significantly, especially when done at scale. Protein powder supplements, commonly derived from leguminous plants including beans and peas, represent cost-effective, accessible sources of concentrated protein and other nutrients that can be used to replace a fraction of the nutrients found in FBS as a minimum. In this study, high-protein powders sourced from marine cyanobacteria (Spirulina), insects (mealworm) and legumes (faba bean) were chosen and assessed for their potential to partially or fully replace serum in media formulations. without compromising the cell viability and growth rate. These sources were selected for their highly nutritionally valuable components, and their potential to be sustainably sourced within the UK food supply chain.

This study was guided by circular bioeconomy principles, which emphasise biological resource reuse, waste minimisation, feedstock and land use efficiency, while respecting ecological boundaries and enhancing societal value (Muscat et al., 2021; Picanço Rodrigues & Fonseca, 2024). Additionally, the selected food-grade components are commonly used as animal feeds. Redirecting agricultural products, such as faba beans, from animal feed toward feedstocks for cultivated meat can provide environmental and economic co-benefits for the food system as an alternative supply chain for feed producers, especially in the event of livestock production decline (Newton & Blaustein-Rejto, 2021).

Insect and mollusks protein isolates have been previously tested for their use in cultivated meat. Batish, Zarei, Nitin, & Ovissipour (2022) assessed protein hydrolysates from insects such as black soldier flies, crickets, and lugworms, and mollusks such as oysters, and mussels, for their suitability for cultivated fish production. All but the cricket protein isolates were able to increase cell growth rates when added to medium with reduced serum down to 2.5% v/v. Kim et al. (2023) also found that by supplementing with edible insect hydrolysates, up to 50% of serum could be replaced for the growth of porcine satellite cells. Mealworms present opportunities for waste minimisation, as they can be raised on organic waste, including agricultural byproducts and food scraps (van Broekhoven, Oonincx, van Huis, & van Loon, 2015).

Plant protein extracts are another example of sustainable protein sources that have already demonstrated utility of replacing some or all the components in serum, while being able to improve cell growth rates (Andreassen, Pedersen, Kristoffersen, & Rønning, 2020; Farges-Haddani et al., 2006). In another study, plant protein extracts were able to replace specific proteins such as albumin (Stout et al., 2023). Faba beans, an underutilised leguminous crop, require minimal fertiliser inputs due their capacity to fix atmospheric nitrogen, reducing nutrient run-off while also enhancing soil health and soil moisture retention through their deep rooting capacities (Multari, Stewart, & Russell, 2015).

Spirulina is a fast-growing marine cyanobacterium which, while requiring minimal land and water resources, holds the capacity to sequester carbon (Shabani, Sayadi, & Rezaei, 2016). In a food context, spirulina (*Arthrospira Platensis)* is a multicellular marine cyanobacterium that has been commonly used as a food supplement for both animals and people. Liestianty et al. (2019) analysed the nutritional content of a spirulina-based powder and found it to have 67% w/w protein, 15% carbohydrates, 0.9% lipids and 1.2% salts, together with a broad range of essential and non-essential amino acids. Spirulina was also found to be a source of gamma linoleic acid, and beta carotene (Choopani, Poorsoltan, Fazilati, Latifi, & Salavati, 2016; Hassaan et al., 2021); both nutritional components that are necessary for cell maintenance in culture. There is precedent for cyanobacteria extracts being used as an alternative to FBS. For example, Jeong et al., (2021) created an extract from *Arthrospira maxima* which was successfully used to replace FBS in the culture of a H460 cancer cell line.

In the context of cultivated meat, the supplementation of culture media with food-grade sources of nutrients has predominantly focused on producing protein hydrolysates which are a mixture of polypeptides, oligopeptides, and amino acids produced from full or partial protein hydrolysis (Cruz et al., 2023). There are clear advantages of breaking down proteins into bioavailable forms for the cells to easily metabolise. However, the processing steps may remove or degrade certain components possibly beneficial to cell growth and viability. As such, the method of pre-processing choice is extremely important for extraction of high-quality nutrients.

This study aimed to investigate food grade protein-rich extracts as partial or full replacements of FBS for cultivated meat production. Additionally, the effects of different methods of processing and different solvents on C2C12 mouse myoblasts and primary porcine myosatellite cell cultivation was investigated.

## Materials and Methods

### Cell culture

Porcine myosatellite cells (pMyoSCs) were isolated using enzymatic dissociation of muscle tissue from a small pig sourced from the Royal Veterinary College (RVC) London following ethical approval. Cells were isolated in flasks coated with Laminin 511 E8 (Merck Millipore; CC160) at a concentration of 0.5 μg/cm². Myogenic potential of the cell population was verified using myogenic differentiation medium comprising DMEM/F12 basal medium supplemented with 2% v/v Horse Serum (Gibco, UK). The C2C12 cell line was also used in this study, and it was sourced from Merck (Merck, 91031101). C2C12 is an immortalised murine myoblast cell line, initially described and isolated by (Yaffe & Saxel, 1977).

Cells were cultured at 37°C with 5% CO₂ in a humidified incubator. C2C12 cells were cultured on uncoated tissue culture treated T-Flasks and microwell plates, while pMyoSCs were cultured on flasks and plates coated with 0.1 μg/cm² laminin-511. Growth Media (GM) for both C2C12 and pMyoSCs comprised of DMEM/F12 (Gibco, UK) basal medium supplemented with 20% v/v heat-inactivated FBS (South America origin, Gibco, UK) and 5 ng/mL human recombinant basic fibroblast growth factor 2 (FGF2) (154aa; Peprotech, UK). During routine culture, cells were passaged at approximately 70% confluency using 1x Tryple Select (Gibco, UK).

### Processing of the selected food grade supplements

Three food grade supplements that fit the criteria of protein-rich, nutritious and sustainable sources were selected and screened for their potential to be used as partial or complete replacements of FBS to support the growth of C2C12 and porcine MyoSCs. The selected food grade supplements are commercially available found in naturist stores **(Table 1)** and they come in a powdered form that is mostly water insoluble. Therefore, these powders had to undergo additional processing prior to their use in cell culture.

**Table 1:** Origin, source, and cost of the food-grade protein concentrates tested.

| Component | Species | Supplier |
| --- | --- | --- |
| Spirulina powder | *Arthrospira platensis* | Naturya, Bath, UK |
| Faba Bean powder | *Vicia faba* | Pulsin, Gloucester, UK |
| Buffalo Mealworm powder | *Alphitobius diaperinus* | Ento Kitchen, Mirfield, UK |

#### Step 1: Initial processing method

The initial method chosen for producing the extracts was ultrasonication. Briefly, the powders were resuspended in the solvent to make 5% w/v (50 g/L) solutions, then vortexed for 5 minutes and sonicated using the Soniprep 150 Sonicator for 6 cycles of 10 seconds each at 23 kHz with the sample tube immersed in ice to ensure no thermal denaturation of the released proteins occurs. Following sonication, the samples were then centrifuged at 1000g for 10 minutes. The pH of the supernatants was then measured and then filtered through a 0.2 μm PES filter.

#### Stage 2: Development of the processing method

The initial solvent used was ultrapure water. However, during the optimisation experiments, the portfolio of solvents used for extraction was extended to also include phosphate buffered saline (PBS) without Calcium or Magnesium (Gibco, 10010023) and 10% v/v vegetable glycerine (Special Ingredients, 5060341114717). While the initial ultrasonication tested used an arbitrary protocol of 6 cycles of 10 seconds each, during the processing optimisation, the number of cycles was increased up to 24, while maintaining the same cycle duration. Additionally, further methods for nutrients extraction were investigated and these included: autoclaving and freeze-thaw cycles. Autoclaving was carried out using a media cycle at 120℃ for 20 minutes. For the freeze-thaw cycles, samples were snap-frozen in dry ice and then immediately thawed out in a water bath at 37℃ for either one or two cycles.

### Testing of food grade protein-rich extracts with cells

#### Initial testing

Initial cytotoxic studies were performed on C2C12 cells. The food grade protein extracts were added to DMEM/F12 with no additional supplementation to achieve concentrations of 0.02, 0.2 and 2% v/v. C2C12 cells were seeded in GM (DMEM/F12 with 20% FBS and 5 ng/mL FGF2 at 5,000 cells/cm² for 24 hours to allow cell attachment before changing to the media formulations containing food grade protein extracts. Cells were cultured in these formulations at 37°C and 5% CO_2_ in a humidified incubator for 4 days, after which, they were harvested and counted. Media formulations were supplemented with a specific volume percentage (% v/v) of this final extract, not the initial powder. This approach allowed us to standardise experiments based on the actual soluble component of the extracts rather than the initial protein concentration that may be subsequently lost during processing.

#### Optimisation testing

Based on the findings of the cytotoxic testing experiment, the concentrations of faba bean, spirulina and mealworm extracts were set to 2% v/v, 2% v/v and 0.5% v/v respectively. The initial formulations, prepared at the specified extract concentrations, were supplemented with 1× Insulin-Transferrin-Selenium (ITS) (Gibco, UK), providing final concentrations of 5.5 µg/mL insulin, 5.5 µg/mL transferrin, and 5 ng/mL selenium, along with 5 ng/mL human recombinant FGF2 (Peprotech, UK). Two controls were used for these experiments: GM containing 10% v/v FBS and DMEM/F12 supplemented with insulin, transferring, selenium and FGF2 (ITSF) at the concentrations described above. To test the effects over long term culture, a multiple consecutive passages approach was taken. Briefly, the cells were seeded at 500 cells/cm² in GM for 24 h and then changed to serum-free media formulations for 5 days before harvesting, with media refreshed every 2 days.

In the second iteration of the long-term passaging screen, to accommodate for lower attachment efficiency in serum-free medium, seeding densities were increased to 2,500 cells/cm^2^ and the cells seeded directly into the serum free media. Cultures were passaged when they reached a minimum of 70% confluency and the culture was continued for 14 days or cultures were terminated if insufficient numbers of live cells were harvested in order to re-seed at the required cell density.

To assess the real effects of the serum-free media formulations, the C2C12 cells were preconditioned to serum-free media through a sequential passage and gradual adaptation strategy as shown in **Table 2**. C2C12 cells were seeded directly into the different media conditions at 2,500 cells/cm² and the media changed after a sample had been taken for PrestoBlue analysis at 24 hours.

**Table 2:**
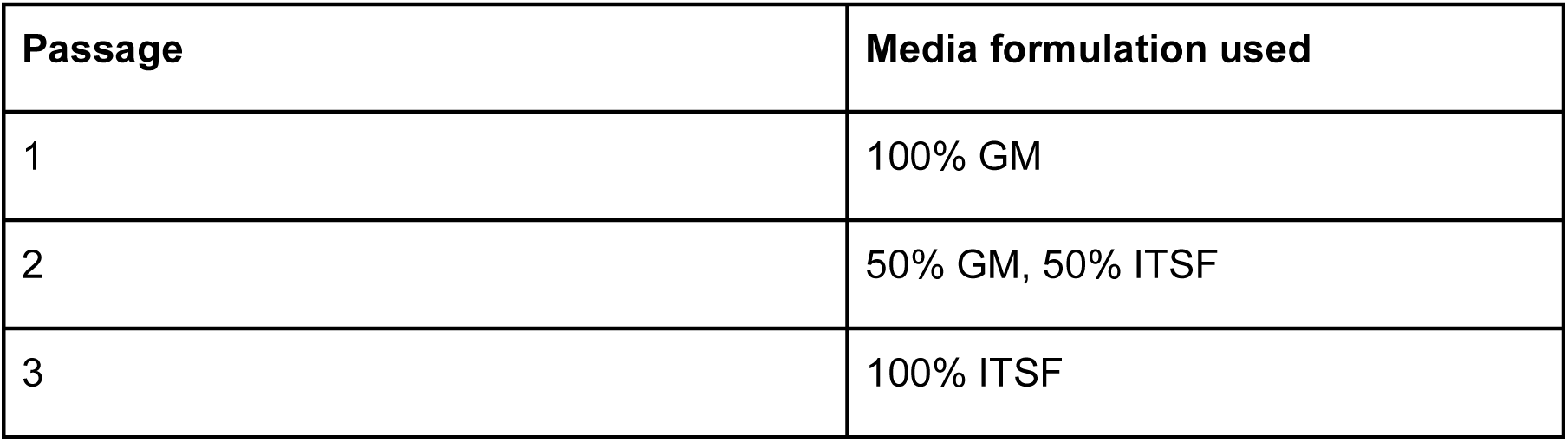
Protocol for gradual serum-free adaptation of C2C12 myoblasts.

### Design of Experiments

To further optimise cell growth over multiple passages in serum-free protein extract formulations, a design of experiment (DoE) approach, full factorial design (2×2×4), was performed to better understand the relationships between the different methods of extraction, food grade extracts and the presence or absence of a selected coating (laminin-511) (**Table 3**). Two different DoE experiments were performed, one for spirulina and one for faba bean extracts supplemented with ITSF. In both experiments, the food grade extracts were obtained through two different methods:

1) <u>method A</u> involving 6 sonication cycles of 10 seconds each, followed by centrifugation at 1000g for 10 minutes;
2) <u>method B</u> involving 12 sonication cycles of 10 seconds each, followed by centrifugation at 5000g for 10 minutes. The food grade extract concentrations ranged from 0.2% 2% to 5% v/v.

**Table 3:** Design of Experiments (DoE) matrix for testing food grade extracts-based media formulations in the culture of C2C12 myoblasts.

| Coating | Extraction method | Food grade extract concentration (% v/v) |
| --- | --- | --- |
| Laminin-511 (0.1µg/cm<sup>2</sup>) | Method A | 0.2 |
| None | Method B | 1.4 |
|  |  | 3.6 |
|  |  | 5 |

There was a total of 32 experimental conditions tested alongside the ITSF-only (no food grade extract supplementation) and GM controls. Both cell attachment and subsequent cell growth were assessed using a Presto Blue assay performed 24 hours and 72 hours post-seeding. Initial models included all main effects, quadratic terms (where applicable). A stepwise backward elimination approach to refine the model was also applied, systematically removing the least significant term (highest p-value > 0.05) in each iteration, while monitoring the R^2^ and the R^2-^adjusted values. The elimination process continued until further term removal degraded the model predictive power.

### Further serum-free media optimisation using the selected food grade extracts

From previous experiments, spirulina was deemed to be the most promising food grade protein-rich extract, and it was taken forward for the further optimisation experiments. A further development involved using a published formulation (B8) as the basal medium (Stout, Mirliani, et al., 2022). B8 media comprised of DMEM/F12 basal medium (Gibco 11320033) with 200 µg/mL L-ascorbic acid 2-phosphate (Sigma-Aldrich 49752), 0.5 µg/mL human recombinant Insulin (Sigma-Aldrich I0908), 0.5 µg/mL human recombinant transferrin (Sigma-Aldrich T3705), 40 ng/mL human recombinant FGF2 (Peprotech, 100-18B), 20 ng/mL sodium selenite (Sigma-Aldrich S5261-10G), 0.1 ng/mL human recombinant NRG1 (MedChemExpress HY-P7365), and 0.1 ng/mL human recombinant TGF-beta 1 (Peprotech, 100-21). B8 was supplemented with the food grade spirulina extracts generated through different processing methods (**Table 4)**. The extracts were added to B8 at 0.002, 0.02, 0.2 and 2% v/v concentrations. pMyoSCs (P3) and C2C12s (P23) were seeded at 2,500 cells/cm² in GM for 24 hours to allow cell attachment. After this, the media was changed with the B8 spirulina extract mixtures. Media was then replaced after 48 hours and assayed 24 hours later using either Presto Blue assay or nuclei quantification with Hoechst 33342. For long term studies, the cultures were kept for 3 consecutive passages with cells passaged every 7 days and media refreshed every 2 days.

**Table 4.** Effect of extraction solvent and cell disruption method on the pH of the spirulina extracts. The mean pH values of 3 measurements are shown.

| Solvent | Processing method | Processing parameter s | Mean pH |
| --- | --- | --- | --- |
| Water | Sonication | 12 cycles of 10s at 23kHz | 6.44 |
| Water | Sonication | 18 cycles of 10s at 23kHz | 6.54 |
| Water | Sonication | 24 cycles of 10s at 23kHz | 6.53 |
| Water | Freeze-thawing | control | 6.63 |
| Water | Freeze-thawing | 1 cycle 30 mins freeze, 45 mins thaw | 6.66 |
| Water | Freeze-thawing | 2 cycle 30 mins freeze, 45 mins thaw | 6.64 |
| Water | Autoclaving | control | 6.55 |
| Water | Autoclaving | 1 cycle | 6.34 |
| PBS | Sonication | 12 cycles of 10s at 23kHz | 6.25 |
| PBS | Sonication | 18 cycles of 10s at 23kHz | 6.23 |
| PBS | Sonication | 24 cycles of 10s at 23kHz | 6.24 |
| PBS | Freeze-thawing | control | 6.47 |
| PBS | Freeze-thawing | 1 cycle 30 mins freeze, 45 mins thaw | 6.50 |
| PBS | Freeze-thawing | 2 cycle 30 mins freeze, 45 mins thaw | 6.49 |
| PBS | Autoclaving | control | 6.47 |
| PBS | Autoclaving | 1 cycle | 6.30 |
| 10% v/v Glycerine/ water | Sonication | 12 cycles of 10s at 23kHz | 6.57 |
| 10% v/v Glycerine/ water | Sonication | 18 cycles of 10s at 23kHz | 6.50 |
| 10% v/v Glycerine/ water | Sonication | 24 cycles of 10s at 23kHz | 6.52 |
| 10% v/v Glycerine/ water | Freeze-thawing | control | 6.63 |
| 10% v/v Glycerine/ water | Freeze-thawing | 1 cycle 30 mins freeze, 45 mins thaw | 6.66 |
| 10% v/v Glycerine/ water | Freeze-thawing | 2 cycle 30 mins freeze, 45 mins thaw | 6.66 |
| 10% v/v Glycerine/ water | Autoclaving | control | 6.55 |
| 10% v/v Glycerine/ water | Autoclaving | 1 cycle | 6.3 |

Based on cell counts, the following parameters were calculated (Hanga et al, 2020):

1. Doubling time (t_d_)

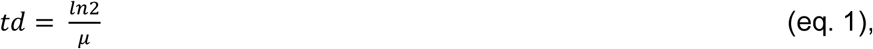
2. Cumulative population doubling level (CPDL)

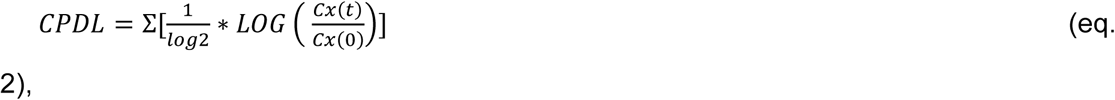

where *Cx(t)* and *Cx(0)* represent cell numbers at the end and start of the culture; t represents time in culture (h).

### Analytical techniques

#### Cell counting

Cell counting and viability were assessed using an NC-3000 automated cell counter using the Via1 cassettes (Chemometec, Denmark). The Via-1 cassettes are preloaded with acridine orange (green, live cells) and DAPI (blue, nuclei) stains. The cell suspension sample is loaded onto the Via-1 cassette where it mixes with the two preloaded dyes. Four images of each quadrant are then taken by the counter in the different fluorescent channels (green and blue). The NC3000 then uses an automated software to count the stained nuclei.

#### Imaging

Cell imaging for routine culture was done on an EVOS XL phase contrast microscope (Thermofisher, UK). For nuclei quantification, cells were stained with 4 µM Hoechst 33342 (Thermo Scientific) for 10 minutes before imaging on the Opera Phenix High throughput cell imager. Images of the whole well were taken using the 5x air objective in non-confocal mode using the Hoechst 333342 channel with 100 ms excitation. Nuclei counts were obtained using the Columbus software and the “count nuclei” protocol.

#### Cell growth assessment

Cell growth assessment was performed using the Presto Blue assay (Invitrogen, A13262, Thermofisher, UK) according to the manufacturer’s protocol. Briefly, spent media was aspirated and a 10% v/v solution of Presto Blue in media was added to the cells and incubated for 1 hour at 37°C in a 5% CO_2_ incubator, wrapped in aluminium foil. Post-incubation, fluorescence measurements were taken at 600 nm using a BMG Labtech Clariostar® microplate reader in black-walled, clear-bottomed tissue culture plates (Thermofisher, 165305). A larger difference in fluorescence indicated a greater amount of reduction from resazurin to resorufin indicating cell growth. A 100% reduced form was produced by autoclaving the dye and it was used as a positive control, while the non-reduced, freshly prepared 10% v/v Presto blue solution was used as a negative control. The percentage reduction of Presto Blue was calculated using equation (3).

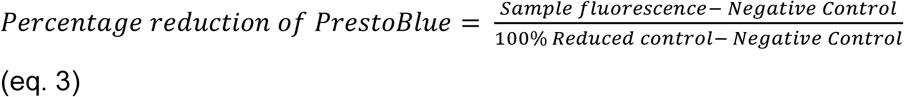

#### FT-IR analysis

An FT-IR analysis was performed to investigate the chemical differences of the food grade extracts. The FT-IR spectra were recorded on a Perkin Elmer Spectrum 2 spectrometer in the spectral range of 4000-400 cm^-^¹ at a resolution of 32 flashes per measurement. Peaks in the FT-IR spectra were identified using the functional data explorer automatic peak identification feature in JMP Pro 18.

#### Amino acid analysis

Amino acid analysis was conducted by Alta Bioscience (Redditch, UK). Samples were deproteinated by mixing 1:1 with a 5% trichloroacetic acid solution, followed by centrifugation and filtration. They were then injected onto a Biochrom 30+ analyser, using ion exchange chromatography with post-column ninhydrin derivatisation and photometric detection.

#### Protein quantification

Protein concentrations in the food grade extracts were measured using the Pierce Bradford assay kit (Thermofisher, 23200) following the manufacturer’s protocol. Briefly, a standard curve was made using a Bovine Serum Albumin standard solution at known concentrations of 0, 2.5, 5, 10, 15, 20 and 25 µg/mL. 5 µl of samples and standards were combined with 250 µl of Bradford assay reagent, shaken for 30 seconds then incubated for 10 minutes at room temperature. The absorbance was then measured at 595nm using the BMG Labtech Clariostar® microplate reader and the protein concentrations were determined from the standard curve.

#### pH and osmolality measurements

The pH was measured with a Mettler Toledo Seven Easy pH meter. Adjustments to pH 7.2 were made by adding 5M NaOH or HCl dropwise until the target pH was reached. Osmolality was measured using an Osmomat 3000 freezing point osmometer.

### Statistical Analysis

All statistical analysis was conducted using JMP 17 and GraphPad Prism 10.3.1. Design of Experiments experimental designs were created using JMP 17. For short-term cell growth studies, an ANOVA was used with a Dunnett’s post hoc test between the media conditions and the control. The tests would only indicate differences at the timepoint of cell counting. Data is shown as Mean ±SD. Three replicates were performed for all conditions tested.

## Results and Discussion

### Cell culture screening of ultrasonicated protein powders

To test the suitability of the selected food grade extracts for cell culture, the extracts obtained using the initial processing method were added at different concentrations to DMEM/F12 alone with no FBS or other supplements. The media formulations produced were then tested with C2C12 cells in a 4-day culture. A significant increase in growth (**p<0.001) was obtained in the 2% v/v faba bean extract-containing formulation. Conversely, the mealworm extract at the highest concentration tested (2% v/v) had a cytotoxic effect resulting in a significant drop in the cell number and viability (****p<0.0001) (**Figures 1A and 1B**).

**Figure 1:**
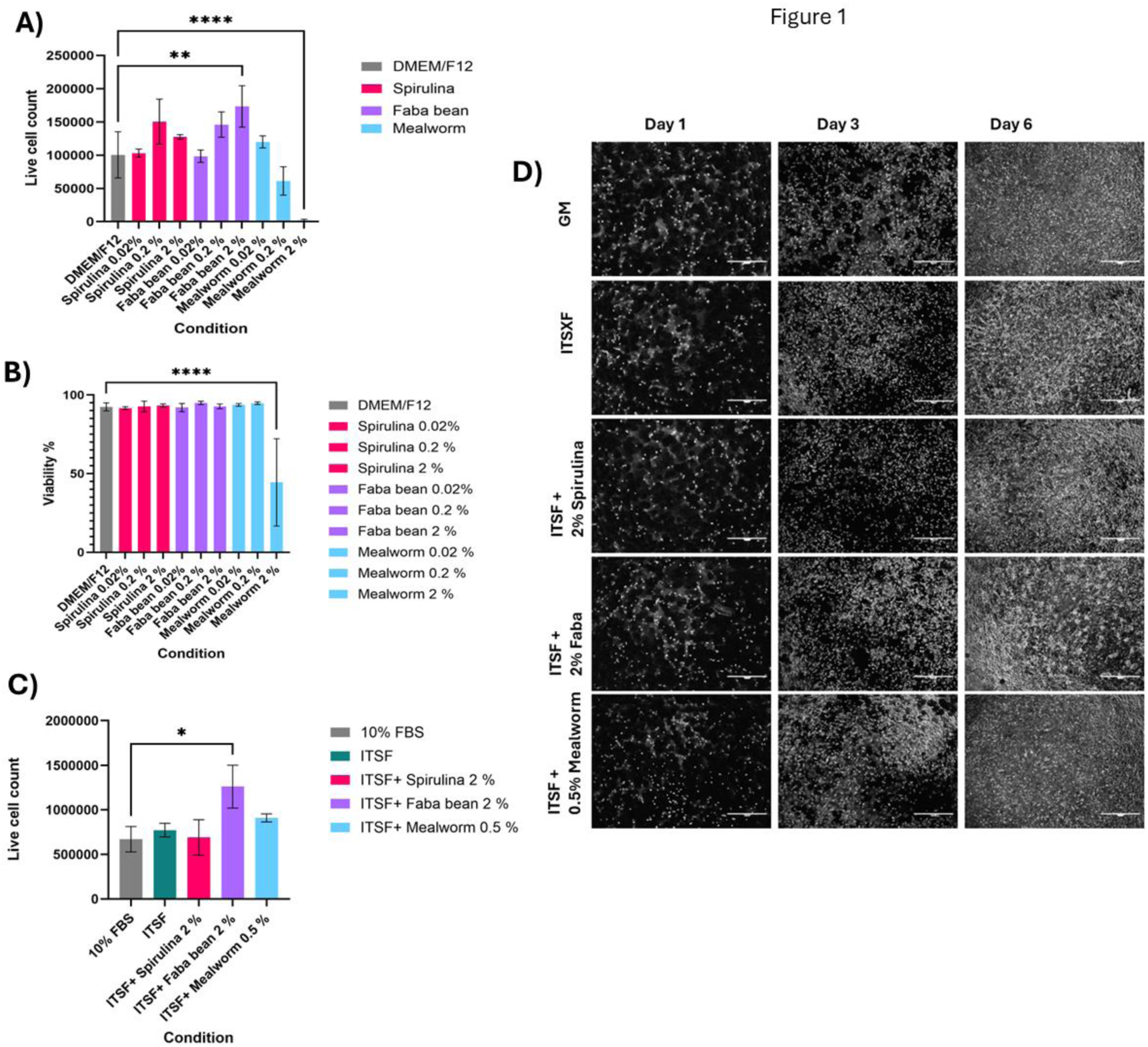
[A-B] C2C12 cells seeded at 5000 cells/cm² cultured for 4 days in food grade extract formulations at different concentrations. A) Live cell count. Data is shown as Mean +/- SD (n=3). B) Cell viability. Data shown as Mean +/- SD (n=3). **P ≤ 0.01; ****P ≤ 0.0001. [C-D] C2C12 cells seeded at 500 cells/cm² at day 6 when cultured in ITSF formulations supplemented with 2% v/v faba bean or spirulina extracts. Two controls were used: GM and ITSF (no supplementation). C) Live cell count. Data shown as Mean ± SD (n=3); *p<0.05. D) Background subtracted and enhanced phase contrast images. Scale bars represent 1000 µm. Background extraction was executed using rolling ball correction and enhanced image after sequential application of contrast stretching (10-90 percentile), gamma correction (γ = 0.5), and CLAHE.

The next formulations tested included supplementation with some essential components for cell growth (insulin – transferring - selenium - FGF2; ITSF) and with the best performing concentrations of food grade extracts. Due to the observed toxicity, mealworm extracts were excluded from further study. From the tested formulations, ITSF with 2% v/v faba bean extract surpassed the cell growth of the serum-based control within the 4-day timeframe (*p<0.05) **(Figure 1C).** Phase contrast images showed the influences of the different media formulations on cell growth and morphology over six days of culture. Phase contrast imaging also revealed that in the ITSF+ 2% v/v faba bean formulation induced cell aggregation **(Figure 1D)**.

The food grade protein extracts were then tested over multiple consecutive passages which is essential to ensure that any residual effects of FBS are removed, while testing if similar growth rates can be maintained longer-term. **Figure 2A** shows similar population doubling levels to the controls for C2C12 cells cultured in faba bean extract-containing formulations and the ITSF control for the first passage at 5 days, whereas a significantly lower (**p<0.01) population doubling level was observed in the ITSF. In contrast (**Figure 2B)**, in the subsequent passage (a further 5 days), there was no cell growth in either of the food grade extracts formulations when no coating was used (****p<0.0001). When using a laminin-511 coating in the second subsequent passage, some cell growth was achieved in the 2% v/v faba bean extract formulation (**Figure 2C**) however still significantly lower than the 10% FBS control (**p<0.01). No growth was achieved in the 2% v/v spirulina extract formulation regardless if laminin-511 was used or not (**Figures 2B and 2C**).

**Figure 2.**
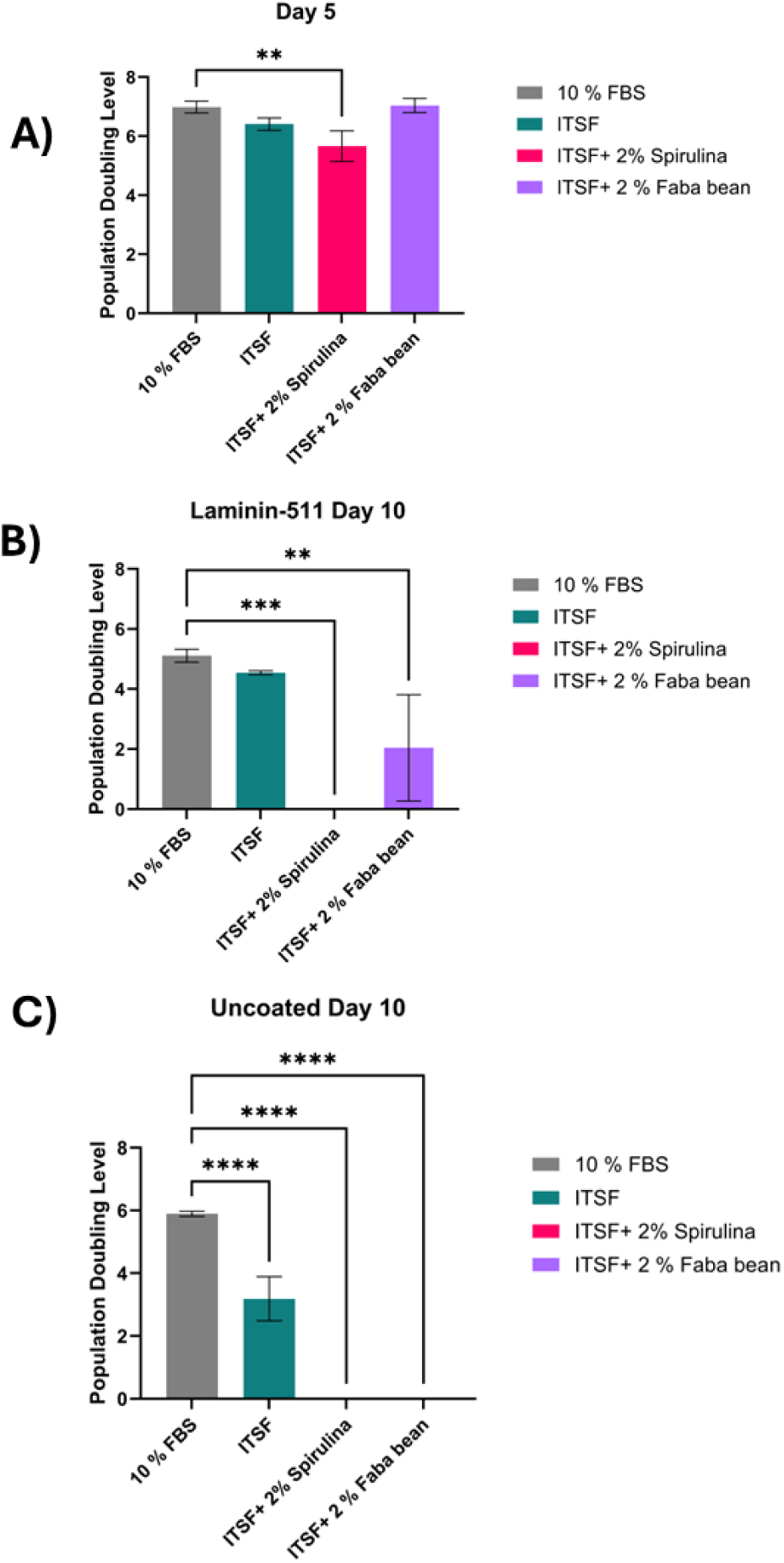
Population doubling levels of C2C12 cells cultured in serum-free media formulations assessed via multi-passage assay. C2C12 myoblasts were seeded at 2,500 cells/cm² in uncoated well plates under serum-free conditions. A) After 5 days, cells were passaged and replated onto either (B) fresh uncoated plates or (C) laminin-511-coated plates, followed by an additional 5-day culture period. Data presented as mean ± SD (n = 3). Statistical significance was evaluated by one-way ANOVA followed by Dunnett’s post hoc test, comparing formulations to the GM control (10% FBS).

C2C12 cells were initially gradually adapted to serum-free medium (schedule shown in **Table 2**) over 3 passages prior to seeding in the DoE approach looking at a series of relevant factors from coating, method of extraction and concentrations of the food grade extract **(Table 3).** When investigating the spirulina extract, the concentration was the only parameter with a significant effect (**** p<0.0001), with the best performance recorded for the 0.2% v/v (**Figures 3A and 3B)**. Optimisation profiling (**Figure 3C)** suggested that the processing method A, with laminin coating and a concentration of 0.02 % v/v spirulina extract was optimal when considering both day one and day 3 performance. None of the conditions tested were significantly different from the ITSF only control at day 1, however the predicted optimal condition at day 3 was significantly greater than the ITSF control (**p< 0.05).

**Figure 3.**
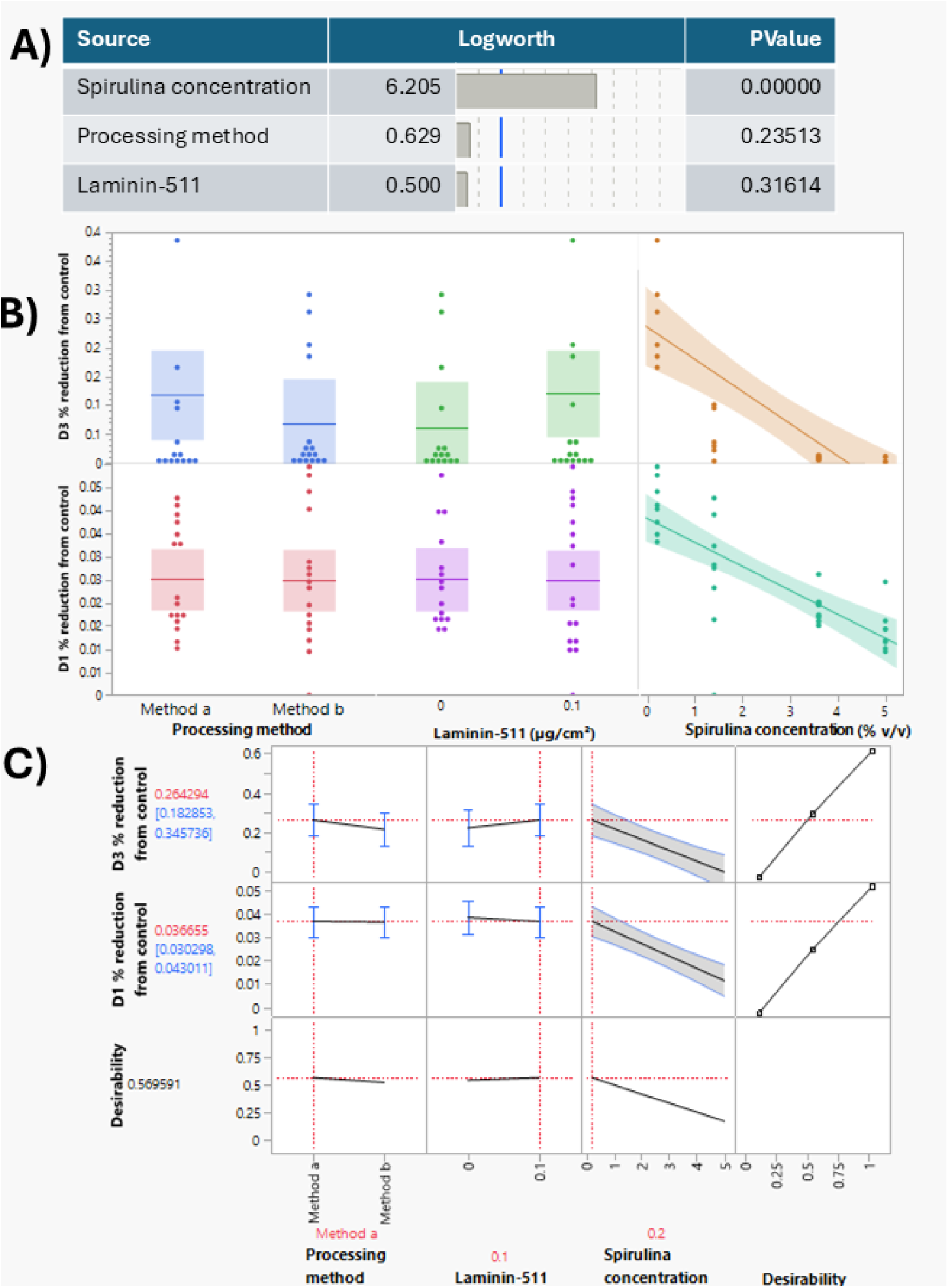
DoE screen investigating Spirulina food grade extract. A) Model effect summary plot; dotted line = logworth of 2 which signals a P value of 0.1. B) Model effect summary plot; growth data normalised to GM control. C) Main effects plot, showing all data points faceted by processing method, laminin coating and spirulina extract concentration % v/v data shown as mean with confidence band, growth data normalised to GM control.

When investigating the faba bean protein extract, the concentration was also found to be a significant factor (**p<0.01) (**Figure 4)**. With slightly better performance at day 1, the processing method B was identified as the optimal processing parameter when used in conjunction with the laminin-511 coating and the lowest concentration of 0.2% v/v. However, this was not significantly different from the ITSF control at day 3 (p>0.05), suggesting that the supplementation of faba bean extracts in the media formulations conferred no cell growth advantages.

**Figure 4.**
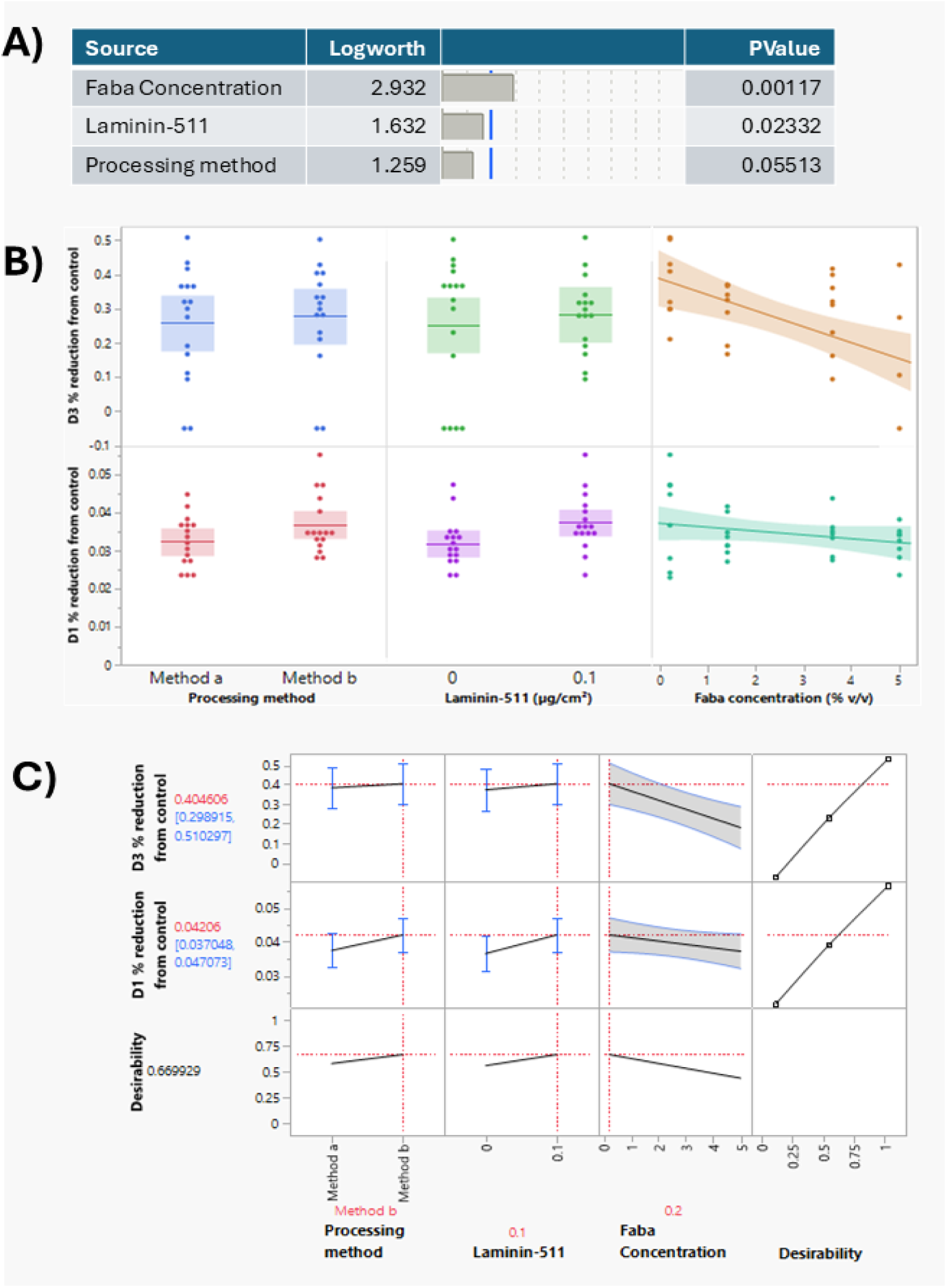
DoE screen investigating Faba bean food grade extract. A) Prediction profiler shows optimal conditions to maximise desirability across day 1 and day 3. B) Main effects plot, showing all data points faceted by processing method, laminin coating and spirulina extract concentration % v/v data shown as mean with confidence bands, growth data normalised to GM control. C) Prediction profiler shows optimal conditions to maximise desirability across day 1 and day 3, growth data normalised to GM control.

Based on the results from the DoE experiments, spirulina extract with its corresponding optimal set of parameters were selected for long term cell culture testing over multiple passages during 14 days of culture. These included 2 candidate media formulations containing 2.6% v/v and 0.2% v/v spirulina produced using Method A (**Figure 5**). Both the ITSF only control and the 0.2% v/v spirulina formulation did not yield enough cells to continue culture at the second passage, leading to culture termination at day 10. Only the GM control and 2.6% v/v spirulina formulation sustained culture for the full 14-day period, with GM reaching a mean cumulative population doublings of 20.54 in that period, while the 2.6% v/v spirulina formulation yielded only 10.73 doublings in the same period (**Figure 5**).

**Figure 5.**
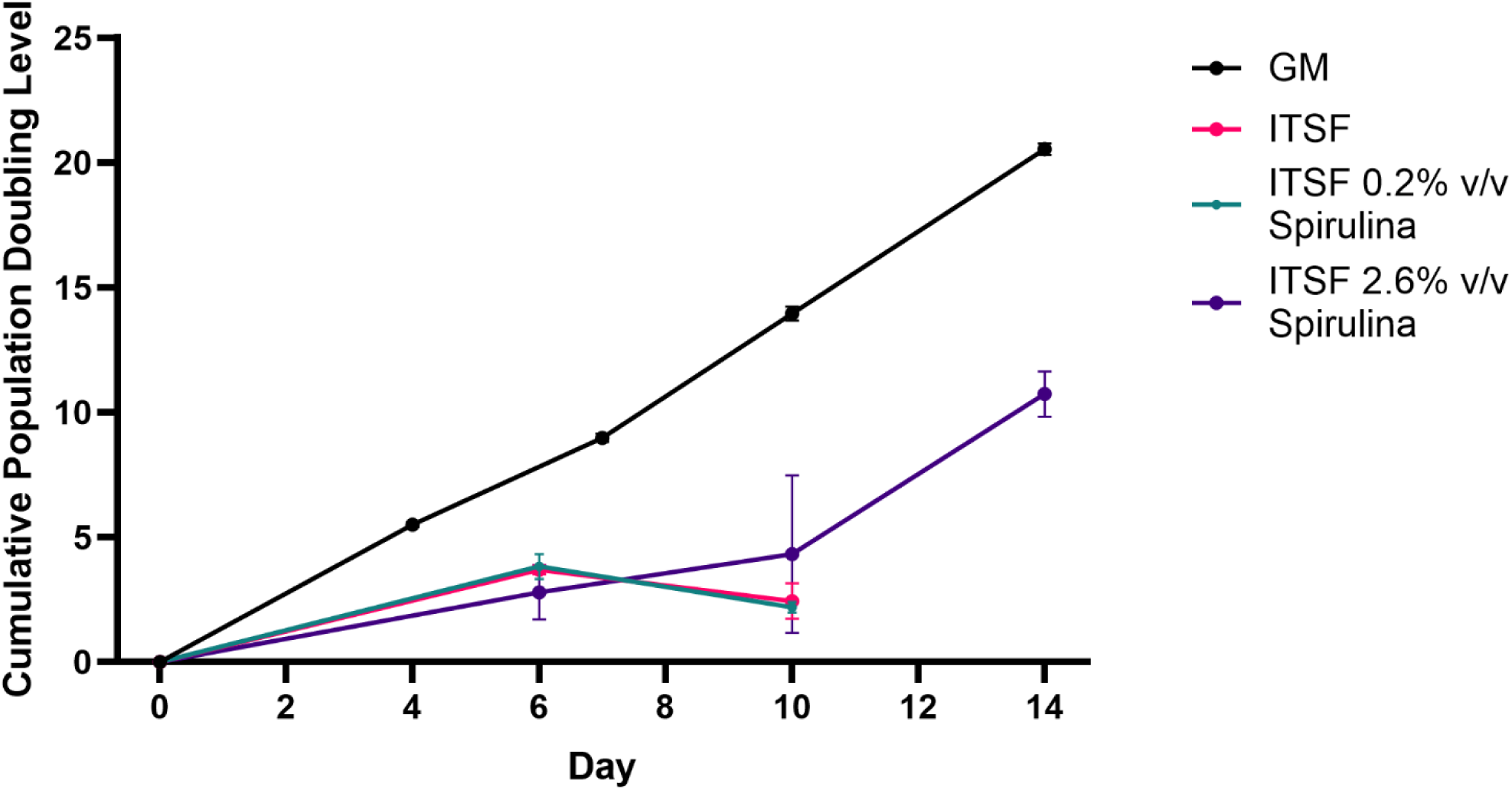
Multi-passage assay of C2C12 cells seeded at 2,500 cells/cm^2^ and passaged at approximately 70% confluency. Cells treated with candidate media formulations suggested from DoE experiments. X indicates not enough cells at passage to re-seed. Data shown as Mean ± SD (n = 3).

The 2.6% v/v spirulina formulation was then selected to be tested with primary pMyoSCs, a more relevant cell type for cultivated meat production. However, after 3 days of culture, the cells detached (**Supplementary Figure S1)**. Because the promising long term growth potential of the spirulina extract did not translate to a more relevant cell type, further optimisation of the processing, media formulation and concentration was performed with the aim of making it effective for the culture of primary cells.

### Spirulina - processing optimisation and composition analysis

#### Processing optimisation

The processing of spirulina extracts was further optimised to improve cell growth with both C2C12 and pMyoSC cells. In the initial processing studies, only one solvent (water) was used. To further optimise the extraction method, three extraction solvents were identified and investigated. These included: water, PBS and 10% v/v vegetable glycerine-water. The glycerine-water system is a food grade solvent with potential for enhanced extraction of compounds from plants (Kowalska, Baj, Kowalski, & Szymańska, 2021). The number of sonication cycles was also increased from 6 and 12 in the initial processing to 18 and 24 cycles of the same duration each (10 s). Several other processing methods for cell lysis to release the nutrients were also investigated and these included 1 and 2 freeze-thaw cycles and autoclaving as a form of heat and pressure treatment which is more likely to hydrolyse proteins into peptides and amino acids.

To assess the efficiency of these changes in the processing methods, protein concentration was measured for all conditions. The glycerine-water solvent when used with 18 or 24 cycles of sonication produced the highest protein concentration, however with no significant difference between the number of cycles (mean 11,478.8 µg/ml and 11,448.7 µg/ml vs 3,036.5 µg/ml in the control). Autoclaving and freeze thawing yielded the lowest protein concentrations regardless of the solvents used, potentially due to protein degradation as a result of exposure to high/low temperature and pressure during processing. The glycerine/water extraction resulted in higher protein concentrations, potentially due to this solvent’s properties of enhanced protein stabilisation and reduced aggregation (Grudniewska et al., 2018; Vagenende, Yap, & Trout, 2009) (**Figure 6A).**

**Figure 6:**
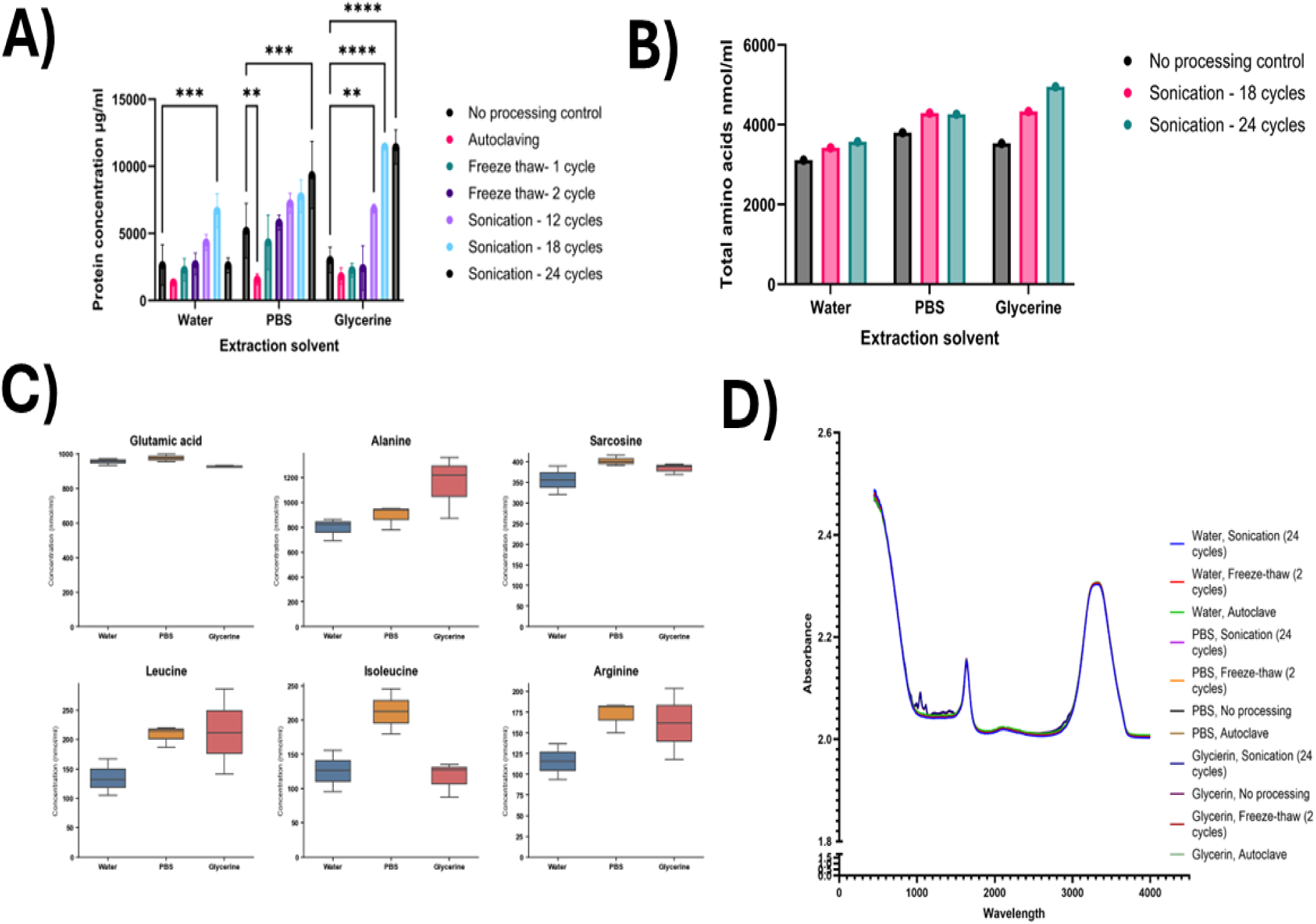
Spirulina extract analysis. A) Protein concentration in spirulina extracts as influenced by the processing methods and solvents. Protein concentrations (µg/mL) were measured in pure spirulina extracts obtained using different processing methods and solvent systems. Values represent mean ± SD (n = 3). Statistical significance was assessed using one-way ANOVA followed by Dunnett’s post hoc test, comparing each treatment group to the unprocessed control (*P ≤ 0.05, **P ≤ 0.01, ***P ≤ 0.001, ****P ≤ 0.0001). B) Total free amino acids analysis of spirulina extracts obtained by sonication (18 and 24 cycles) when using different solvents. C) Total free amino acids of top 6 amino acids of sonication cycles 18 and 24 cycles D) FT-IR analysis spectra of spirulina extracts.

Amino acids profiles were also determined for the conditions that yielded the highest protein concentrations (**Figure 6B**). The glycerine/water solvent when paired with 24 sonication cycles, yielded the highest total amino acids concentration of 4950 µM compared to 4260 µM in PBS and 3570 µM in water. This condition also had the highest yield of essential amino acids. Compared to no processing controls, sonication yielded higher amino acid concentrations in all conditions tested. **Figure 6C** shows the relative concentrations of the 6 most abundant free amino acids (*e.g*. glutamic acid, glycine, alanine, leucine, isoleucine and arginine). These amino acids are typically prevalent in Spirulina (Clément, Giddey, & Menzi, 1967). The optimised formulation resulted in a 4950 µM concentration of free amino acids. However, when diluted to its effective concentration (0.2-2.6% v/v), this equals to only 9.9–128.7 µM. This concentration is very low compared to the 7.25 mM total amino acids already present in the DMEM/F12 basal medium. Therefore, the contribution of free amino acids of the spirulina extract obtained was likely to have a negligible effect on the cells in these serum-free media formulations.

To get a further understanding of the potential components found in the spirulina extracts produced using different methods of processing and solvents, Fourier-transform infrared (FT-IR) was used (**Figure 6D; Supplementary Table S1).** All tested spirulina extracts exhibited peaks around 3306 cm^-1^, 2119 cm^-1^, 1631 cm^-1^ and 450 cm^-1^. The peaks in the 1935-1640 cm^-1^ region indicate Amide 1 C=O vibration, typically seen in carbonyl groups which can be found in proteins, lipids carbohydrates or secondary metabolites (Liu, Zhu, Li, & Zhou, 2022). The broad peak observed in the 3300-3400 cm⁻¹ region is characteristic of O-H or N-H stretching vibrations. Its broad shape is a direct result of hydrogen bonding between the molecules, which creates a range of slightly different bond energies and thus a wide absorption band (Coates, 2000). Processing differences were observed. For example, the spirulina extracts produced using sonication showed a broadening of peaks at 1635-1640 cm^-1^ compared to the extracts produced using freeze-thaw and the control groups, indicating structural heterogeneity. Differences in the spectra also emerged from the samples extracted with 10% v/v glycerine/water solvent. These included additional peaks at 1043 and 1112 cm^-1^ which are likely from the C-OH of primary and secondary alcohol of glycerol (Yan, Li, & Zhang, 2017). The FTIR spectra indicates that the spirulina extracts might contain other components such as lipids alongside proteins and amino acids which are essential for cell maintenance.

Other considerations when developing serum-free media formulations to avoid potential cytotoxicity include suboptimal pH and osmolality. pH out of range of the physiological optimum of 7.4 may cause alterations in enzyme activity, protein folding and DNA stability (Chaudhry, Bowen, & Piret, 2009; Gillies, Martinez-Zaguilan, Peterson, & Perona, 1992). Osmolality also needs to be tightly regulated to maintain cell volume and a stable internal cellular environment (Chaudhry et al., 2009; Michl, Park, & Swietach, 2019). The pH values of the spirulina extracts were measured post-processing and are shown in **Table 4**. All extracts were slightly acidic. The autoclaved sample had the lowest pH, most likely because the process hydrolysed proteins, releasing acidic amino acids that increased the hydrogen ion (H⁺) concentration. However, after resuspension of the extracts at the targeted concentration in the B8 formulation, the pH and osmolality of all tested extracts were within the optimal range of 7.2 to 7.4 and 280-320 mOsm/kg respectively.

#### Growth analysis of optimised extractions

The optimised spirulina extracts were then tested at a concentration range of 0.002-2% v/v on both C2C12 myoblasts and pMyoSCs cells, however this time in a more complex formulation using the B8 serum-free media formulation previously published (Kuo et al., 2020) as the base. The cells were seeded in GM for 24 hours and then the medium was changed to the enhanced spirulina extract-containing formulations as described above. Cell growth was assessed through nuclei count (Hoechst assay) and viability/metabolic activity through Presto Blue on day 4 of the culture. Notably, the growth rate of the C2C12 myoblasts in the B8 serum free medium was much closer to the GM control with B8 being 64.47% of the mean nuclei count and 99.89% of the mean Presto Blue reduction in GM, whereas for pMyoSCs, B8 was only 28.31% of the nuclei count and 62.52% of the Presto Blue reduction in GM. The variance in results between the Presto Blue and the nuclei count may be due to the Presto Blue assay being an indicator of cellular metabolism rather than direct cell number. When compared, the nuclei count was positively correlated with the Presto Blue response (R² = 0.280 for pMyoSC, R² = 0.003 for C2C12), but with poor predictive accuracy between the two methods (**Supplementary Figure S2)**.

A combined score of both methods, taking the average of the nuclei count and PrestoBlue normalised to the GM control per well was calculated (**Figure 7**). For pMyoSC (**Figure 7A**), no conditions were significantly higher than the B8 control, however 4 conditions were significantly lower than the B8 control combined score of 45.2%. These were: Freeze-thaw PBS 0.02 % w/v, No-processing PBS 2% w/v, Sonication Glycerine 2% w/v, and Sonication PBS 2% w/v. The highest combined score was for pMyoSCs water, sonication, 0.02% w/v with an average of 58.5% difference from the GM control.

**Figure 7:**
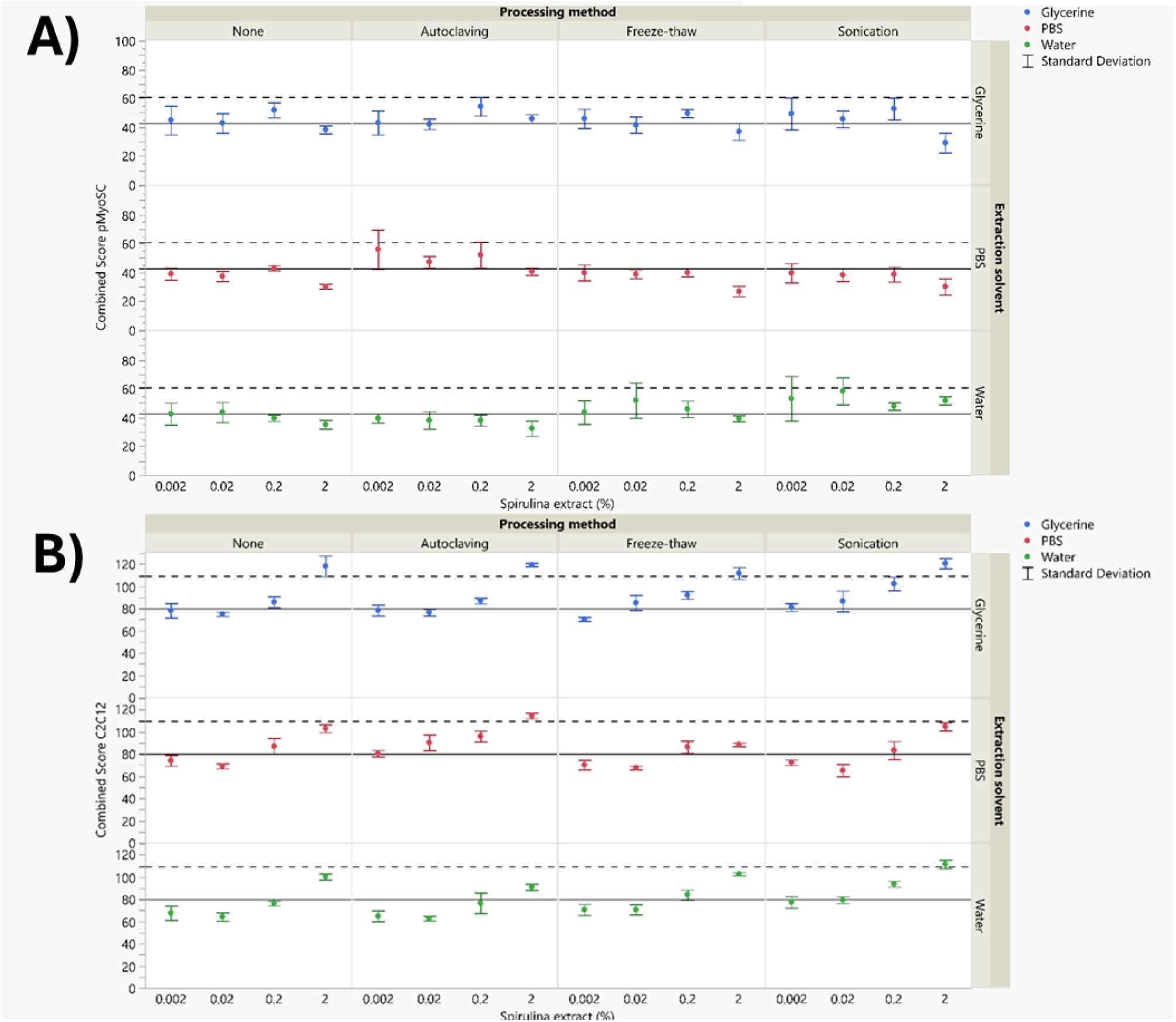
Combined score obtained by averaging the Presto blue data and Hoechst nuclei count per well showing the effect of spirulina extracts at different concentrations obtained using different processing methods and solvents. The solid line indicates the mean difference from the GM control for the B8 only control and the dotted lines indicate the mean difference from the GM control for the B8 only control + 2 standard deviations. A) Tested with pMyoSCs; B) Tested with C2C12 myoblasts.

For C2C12s (**Figure 7B**), there were 13 conditions significantly higher than the B8 control average combined score of 82.2%, with 11 of these surpassing the GM control. The 3 highest values obtained were all using the glycerine/water extraction solvent at a concentration of 2% v/v. Sonication when using glycerine and spirulina extract at 2% v/v resulted in a 120.89% positive difference from the GM control. 9 conditions were significantly lower than the GM control. The method of 24-cycle sonication in a 10% glycerine/water solvent yielded the highest protein concentration, at approximately 11.45 mg/ml and a 20% increase in growth for C2C12s. For context, when this extract was added to the B8 medium at the effective concentration of 2% v/v, it contributed to a final soluble protein concentration of approximately 0.2 g/L to the culture environment.

The 24-cycle sonication condition in PBS at 0.2% v/v spirulina concentration yielded the highest average nuclei count for C2C12 cells at 92.47% of the growth of the GM condition, whereas the highest Presto Blue response (176.4% of the rate compared to the GM control) was yielded from 2% v/v spirulina extract obtained using a 24-cycle sonication in glycerine/water. For pMyoSCs, none of the tested conditions significantly increased the number of nuclei. However, the 24-cycle sonication condition in water at a 0.2% v/v concentration yielded the highest number of nuclei (42.21% of the GM control.) This was also the only condition that was above 2 standard deviations from the B8 control. Despite the minimal increases in the number of nuclei, the Presto Blue assay had the strongest response from the same condition (water, 24-cycle sonication), but for the higher concentration of 2% v/v which was significantly higher than the B8 control 92.5 % (***p=0.001). From all processing methods tested, the sonication yielded the highest protein concentration and led to the highest cell growth rates. The processing method (pMyoSC:*** p<0.001; C2C12:* p<0.05), extraction solvent (pMyoSC:* p<0.001; C2C12:* p<0.05) and spirulina concentration (pMyoSC:*** p<0.001; C2C12:*** p<0.001) all had significant effects on pMyoSC and C2C12 nuclei count.

The screening of extracts was deliberately based on the volume percentage added to the media, not on normalising to protein concentration. This methodological choice is key, as the extracts are prepared from lysed whole spirulina cells and thus contain a complex profile of biomolecules beyond just protein, such as vitamins, minerals, and pigments like phycocyanin (Cirkovic Velickovic & Stanic-Vucinic, 2018; Zhang, Huang, Wang, Wan, & Wang, 2024). Normalising to protein would alter the relative concentration of these non-protein components, thereby masking their potential synergistic or inhibitory effects. By dosing via volume, we directly assessed the bioactivity of the complete extract as a potential supplement. Consequently, future studies are warranted to deconvolute these effects by specifically investigating the role of purified protein fractions versus other extract components.

For example, spirulina is known to also contain phycocyanin and phenolic compounds that act as antioxidants (Cirkovic Velickovic & Stanic-Vucinic, 2018; Zhang, Huang, Wang, Wan, & Wang, 2024). These compounds can bind to proteins and minerals, thus reducing their bioavailability to cells when the extracts are used in cell culture. With high concentrations of proteins, carotenoids, and essential fatty acids, spirulina has a high potential to replace many of the roles of foetal bovine serum. There is already evidence of this effect on mammalian cells with Jeong et al., (2021) finding similar rates of apoptosis, morphology, and metabolite levels in the H460 human lung cancer cell line compared to the FBS containing control. In HS2 keratinocyte cells, Gunes, Tamburaci, Dalay, & Gurhan (2017) found the group treated with a spirulina extract to achieve a higher rate of proliferation than the control group.

### Conclusions

This study highlights the utility of spirulina extracts as partial replacements for FBS in cultivated meat production. Simple processing methods can be adopted to convert protein-rich powders into extracts suitable as cell culture supplements. Faba bean and spirulina extracts boosted short term proliferation of C2C12 cells. However, when cultured over multiple passages in simplified media formulations, a large drop in cell proliferation occurred, which can be related to a reduced cell attachment owing to FBS containing attachment factors not being present once FBS is removed. Optimisation of spirulina extracts to support long-term growth and complete replacement of FBS was achieved. While a formulation was identified that indicated higher growth rates for C2C12 cells, no formulation was found to significantly enhance nuclei count for pMyoSCs indicating potentially different nutrient requirements for the different cell types tested. These findings highlight the challenges to be overcome to create a serum-free media formulation that supports long-term growth of porcine primary cells. As a low-cost ingredient with good supply chains, spirulina has strong potential to contribute to the development of a cost-effective, serum-free culture medium for cultivated meat production. However, further work needs to be done to optimise such a formulation.

Beyond its biological efficacy, the use of food-grade spirulina presents a compelling economic advantage for the scalable production of cultivated meat. Based on our initial processing concentration of 50 g/L leading to a final protein concentration of 0.2 mg/L, the current commercial price of spirulina powder is approximately £2.25 per litre. This represents a very low investment for the significant gains in cell growth observed, particularly in C2C12 myoblasts. Whilst downstream processing, such as sterile filtration, will contribute to the final cost, these steps could be integrated efficiently at scale. For instance, the filtration of the spirulina extract could be coupled with the sterilisation of other media components during large-scale batch preparation, thereby minimising operational overhead. This cost-benefit profile strengthens the potential of spirulina extracts as a viable, low-cost supplement for reducing the overall expense of serum-free media in cultivated meat production.

Future work should focus on the replication of these results on other cell types with lower nutritional requirements such as cultivated seafood where algae may be more representative of the nutritional requirements of the organisms. The analysis of the composition of spirulina extracts through methods such as mass spectrometry to identify any components with potential to limit cell growth to mitigate their effects on cells should also be studied. Further serum-free media formulation development should be carried out on pMyoSCs to improve their growth outcomes in serum-free media, incorporating spirulina extracts.

## Acknowledgements

William Gordon-Petrovskii is grateful to the UCL EPSRC Doctoral Training Program (REF: EP/R513143/1) for the award of a PhD studentship.

It was also funded via an EPSRC manufacturing research hub for a sustainable future grant (REF: EP/X038114/1): Cellular Agriculture Manufacturing Hub (CARMA) https://carmahub.co.uk/.

It was also funded via a Transforming UK Food Systems Strategic Priorities Fund Programme grant (REF: B/V011391/1): UK Food Systems Centre for Doctoral Training (UKFS-CDT) https://foodsystems-cdt.ac.uk/.

## Data availability

The datasets generated during and/or analysed during the current study are available from the corresponding author on reasonable request.

## Ethics Approval

Porcine myosatellite cells (pMyoSCs) were isolated using enzymatic dissociation of muscle tissue from a small pig sourced from the Royal Veterinary College (RVC) London following ethical approval.

## Abbreviations

ANOVA: Analysis of Variance
B8: A defined serum-free basal media formulation (Stout et al., 2022)
BSA: Bovine Serum Albumin
C2C12: An immortalized mouse myoblast cell line
CPDL: Cumulative Population Doubling Level
CLAHE: Contrast Limited Adaptive Histogram Equalization
DMEM/F12: Dulbecco’s Modified Eagle Medium/Nutrient Mixture F-12
DoE: Design of Experiments
ECM: Extracellular Matrix
FBS: Foetal Bovine Serum
FT-IR: Fourier-Transform Infrared Spectroscopy
FGF2: Fibroblast Growth Factor 2
GM: Growth Medium (typically containing FBS)
HCl: Hydrochloric Acid
ITS: Insulin-Transferrin-Selenium
ITSF: Insulin-Transferrin-Selenium-FGF2 (a serum-free formulation)
NaOH: Sodium Hydroxide
NRG1: Neuregulin 1
PBS: Phosphate Buffered Saline
PES: Polyethersulfone
pMyoSCs: Primary Porcine Myosatellite Cells
RVC: Royal Veterinary College
SD: Standard Deviation
SF: Serum-Free
TGF-β1: Transforming Growth Factor Beta 1
v/v: Volume per Volume
w/v: Weight per Volume

