## Supplementary materials for "Investigations into cyanobacteria, plant, and insect protein extracts as serum-replacement supplements for the expansion of cells for cultivated meat"

*Supplementary Figure S1: Phase contrast images of ITSXF+ 2.6% spirulina extract on day 1, 2 and 5. Scale bars represent 1000 μm*

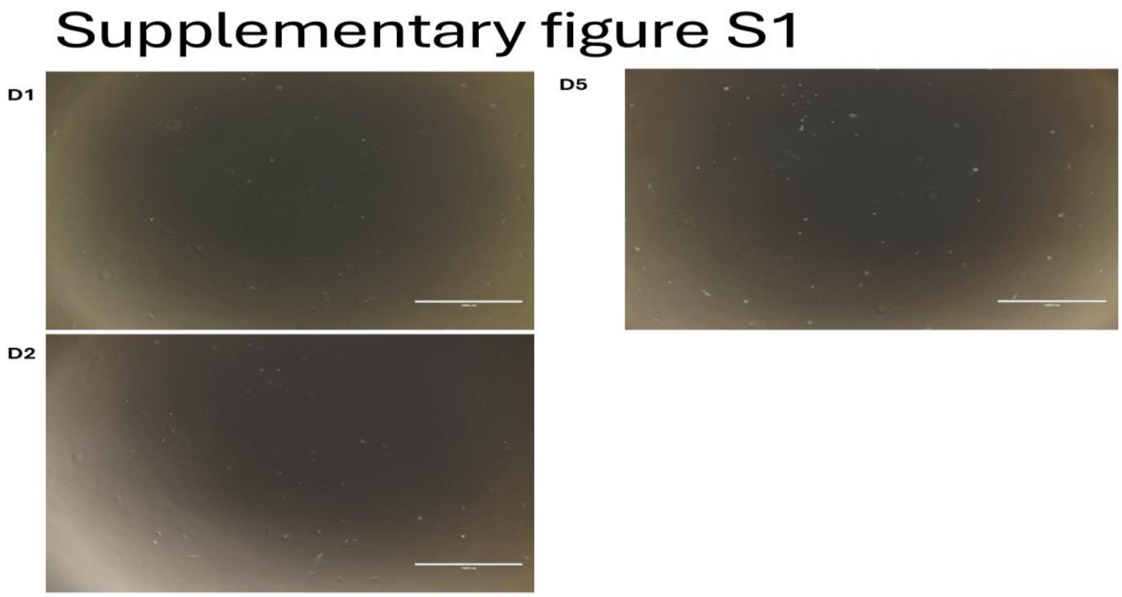


*Supplementary Figure S2: Scatterplots between PrestoBlue and Nuclei Count for a) pMyoSCs and B)C2C12s*

### Supplementary figure 2

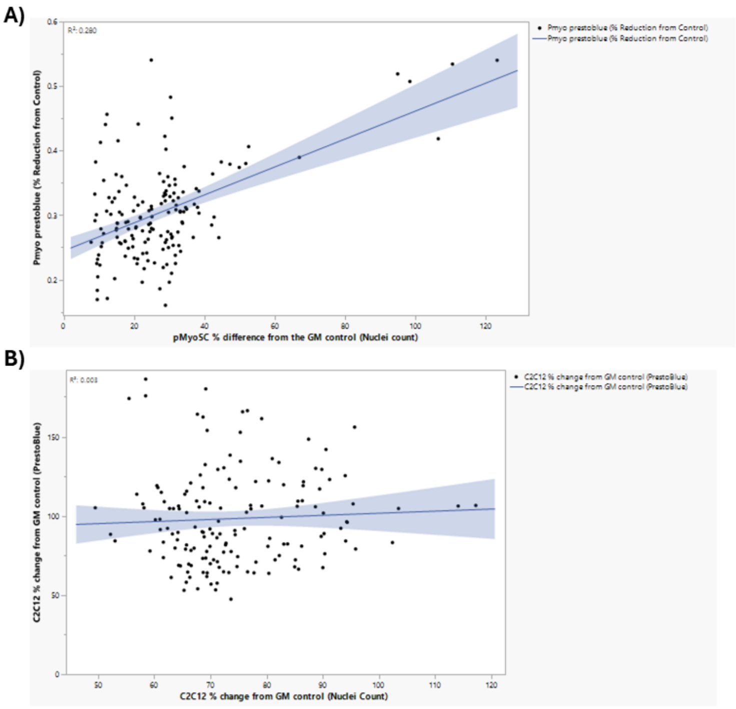


**Table S1, extracts tested in initial growth studies**

*Supplementary Table S1: FTIR absorption peaks for spirulina extracts produced using different methods.*

| Protein Powder | Solvent | Process Parameter | Processing variation | Peak ID | Peak Start | Peak End | Location |
|---|---|---|---|---|---|---|---|
| Spirulina | Water | Sonication | 24 cycles | 0 | 450 | 528 | 452 |
| Spirulina | Water | Sonication | 24 cycles | 1 | 1489 | 1843 | 1635 |

| | | | | | | | |
|---|---|---|---|---|---|---|---|
| Spirulina | Water | Sonication | 24 cycles | 2 | 2013 | 2141 | 2119 |
| Spirulina | Water | Sonication | 24 cycles | 3 | 2602 | 3792 | 3306 |
| Spirulina | Water | Sonication | 24 cycles | 4 | 2602 | 3792 | 3338 |
| Spirulina | Water | Freeze-thawing | 2 cycles | 0 | 450 | 483 | 456 |
| Spirulina | Water | Freeze-thawing | 2 cycles | 1 | 528 | 1065 | 535 |
| Spirulina | Water | Freeze-thawing | 2 cycles | 2 | 1464 | 1842 | 1635 |
| Spirulina | Water | Freeze-thawing | 2 cycles | 3 | 2060 | 2133 | 2119 |
| Spirulina | Water | Freeze-thawing | 2 cycles | 4 | 2060 | 2133 | 2119 |
| Spirulina | Water | Freeze-thawing | 2 cycles | 5 | 2848 | 4000 | 3307 |
| Spirulina | Water | Freeze-thawing | 2 cycles | 6 | 2848 | 4000 | 3307 |
| Spirulina | Water | Freeze-thawing | 2 cycles | 7 | 2848 | 4000 | 3339 |
| Spirulina | Water | Autoclaving | 1 cycle | 0 | 488 | 1061 | 495 |
| Spirulina | Water | Autoclaving | 1 cycle | 1 | 1464 | 1842 | 1635 |
| Spirulina | Water | Autoclaving | 1 cycle | 2 | 1464 | 1842 | 1635 |
| Spirulina | Water | Autoclaving | 1 cycle | 3 | 2028 | 2058 | 2051 |
| Spirulina | Water | Autoclaving | 1 cycle | 4 | 2093 | 2133 | 2112 |
| Spirulina | Water | Autoclaving | 1 cycle | 5 | 2175 | 2209 | 2188 |
| Spirulina | Water | Autoclaving | 1 cycle | 6 | 2848 | 3745 | 3306 |
| Spirulina | Water | Autoclaving | 1 cycle | 7 | 2848 | 3745 | 3338 |
| Spirulina | PBS | Sonication | 24 cycles | 0 | 450 | 483 | 457 |
| Spirulina | PBS | Sonication | 24 cycles | 1 | 1464 | 1844 | 1635 |
| Spirulina | PBS | Sonication | 24 cycles | 2 | 2013 | 2061 | 2050 |
| Spirulina | PBS | Sonication | 24 cycles | 3 | 2061 | 2133 | 2113 |
| Spirulina | PBS | Sonication | 24 cycles | 4 | 2157 | 2174 | 2162 |
| Spirulina | PBS | Sonication | 24 cycles | 5 | 2849 | 4000 | 3307 |

| | | | | | | | |
|---|---|---|---|---|---|---|---|
| Spirulina | PBS | Sonication | 24 cycles | 6 | 2849 | 4000 | 3338 |
| Spirulina | PBS | Freeze-thawing | 2 Cycles | 0 | 450 | 484 | 457 |
| Spirulina | PBS | Freeze-thawing | 2 Cycles | 1 | 1464 | 1842 | 1635 |
| Spirulina | PBS | Freeze-thawing | 2 Cycles | 2 | 2022 | 2061 | 2050 |
| Spirulina | PBS | Freeze-thawing | 2 Cycles | 3 | 2093 | 2134 | 2114 |
| Spirulina | PBS | Freeze-thawing | 2 Cycles | 4 | 2175 | 2210 | 2187 |
| Spirulina | PBS | Freeze-thawing | 2 Cycles | 5 | 2848 | 4000 | 3307 |
| Spirulina | PBS | Freeze-thawing | 2 Cycles | 6 | 2848 | 4000 | 3307 |
| Spirulina | PBS | Freeze-thawing | 2 Cycles | 7 | 2848 | 4000 | 3339 |
| Spirulina | PBS | None | control | 0 | 450 | 484 | 456 |
| Spirulina | PBS | None | control | 1 | 1464 | 1842 | 1635 |
| Spirulina | PBS | None | control | 2 | 2024 | 2060 | 2050 |
| Spirulina | PBS | None | control | 3 | 2060 | 2133 | 2114 |
| Spirulina | PBS | None | control | 4 | 2157 | 2174 | 2163 |
| Spirulina | PBS | None | control | 5 | 2848 | 4000 | 3306 |
| Spirulina | PBS | None | control | 6 | 2848 | 4000 | 3339 |
| Spirulina | PBS | Autoclaving | 1 cycle | 0 | 450 | 484 | 457 |
| Spirulina | PBS | Autoclaving | 1 cycle | 1 | 1488 | 1842 | 1635 |
| Spirulina | PBS | Autoclaving | 1 cycle | 2 | 2024 | 2061 | 2051 |
| Spirulina | PBS | Autoclaving | 1 cycle | 3 | 2061 | 2133 | 2113 |
| Spirulina | PBS | Autoclaving | 1 cycle | 4 | 2174 | 2209 | 2187 |
| Spirulina | PBS | Autoclaving | 1 cycle | 5 | 2849 | 4000 | 3306 |
| Spirulina | PBS | Autoclaving | 1 cycle | 6 | 2849 | 4000 | 3306 |
| Spirulina | PBS | Autoclaving | 1 cycle | 7 | 2849 | 4000 | 3339 |
| Spirulina | Glycerine | Sonication | 24 cycles | 0 | 450 | 506 | 465 |

| | | | | | | | |
|---|---|---|---|---|---|---|---|
| Spirulina | Glycerine | Sonication | 24 cycles | 1 | 980 | 1010 | 994 |
| Spirulina | Glycerine | Sonication | 24 cycles | 2 | 1010 | 1090 | 1043 |
| Spirulina | Glycerine | Sonication | 24 cycles | 3 | 1090 | 1145 | 1112 |
| Spirulina | Glycerine | Sonication | 24 cycles | 4 | 1166 | 1263 | 1212 |
| Spirulina | Glycerine | Sonication | 24 cycles | 5 | 1263 | 1359 | 1336 |
| Spirulina | Glycerine | Sonication | 24 cycles | 6 | 1367 | 1449 | 1411 |
| Spirulina | Glycerine | Sonication | 24 cycles | 7 | 1497 | 1845 | 1639 |
| Spirulina | Glycerine | Sonication | 24 cycles | 8 | 2013 | 2062 | 2051 |
| Spirulina | Glycerine | Sonication | 24 cycles | 9 | 2062 | 2134 | 2114 |
| Spirulina | Glycerine | Sonication | 24 cycles | 10 | 2173 | 2208 | 2187 |
| Spirulina | Glycerine | Sonication | 24 cycles | 11 | 2915 | 4000 | 3306 |
| Spirulina | Glycerine | Sonication | 24 cycles | 12 | 2915 | 4000 | 3338 |
| Spirulina | Glycerine | Freeze-thawing | control | 0 | 450 | 961 | 457 |
| Spirulina | Glycerine | Freeze-thawing | control | 1 | 450 | 961 | 457 |
| Spirulina | Glycerine | Freeze-thawing | control | 2 | 981 | 1011 | 994 |
| Spirulina | Glycerine | Freeze-thawing | control | 3 | 1011 | 1089 | 1043 |
| Spirulina | Glycerine | Freeze-thawing | control | 4 | 1091 | 1159 | 1112 |
| Spirulina | Glycerine | Freeze-thawing | control | 5 | 1166 | 1262 | 1211 |
| Spirulina | Glycerine | Freeze-thawing | control | 6 | 1303 | 1363 | 1336 |
| Spirulina | Glycerine | Freeze-thawing | control | 7 | 1376 | 1450 | 1411 |
| Spirulina | Glycerine | Freeze-thawing | control | 8 | 1494 | 1845 | 1639 |
| Spirulina | Glycerine | Freeze-thawing | control | 9 | 2024 | 2061 | 2051 |
| Spirulina | Glycerine | Freeze-thawing | control | 10 | 2093 | 2133 | 2113 |
| Spirulina | Glycerine | Freeze-thawing | control | 11 | 2174 | 2208 | 2187 |
| Spirulina | Glycerine | Freeze-thawing | control | 12 | 2914 | 4000 | 3306 |

| | | | | | | | |
|---|---|---|---|---|---|---|---|
| Spirulina | Glycerine | Freeze-thawing | control | 13 | 2914 | 4000 | 3338 |
| Spirulina | Glycerine | None | control | 0 | 450 | 963 | 457 |
| Spirulina | Glycerine | None | control | 1 | 450 | 963 | 457 |
| Spirulina | Glycerine | None | control | 2 | 980 | 1011 | 994 |
| Spirulina | Glycerine | None | control | 3 | 1011 | 1089 | 1044 |
| Spirulina | Glycerine | None | control | 4 | 1090 | 1148 | 1112 |
| Spirulina | Glycerine | None | control | 5 | 1168 | 1261 | 1211 |
| Spirulina | Glycerine | None | control | 6 | 1261 | 1365 | 1336 |
| Spirulina | Glycerine | None | control | 7 | 1378 | 1450 | 1412 |
| Spirulina | Glycerine | None | control | 8 | 1498 | 1841 | 1638 |
| Spirulina | Glycerine | None | control | 9 | 1498 | 1841 | 1638 |
| Spirulina | Glycerine | None | control | 10 | 2021 | 2061 | 2050 |
| Spirulina | Glycerine | None | control | 11 | 2093 | 2133 | 2115 |
| Spirulina | Glycerine | None | control | 12 | 2174 | 2217 | 2187 |
| Spirulina | Glycerine | None | control | 13 | 2914 | 3744 | 3306 |
| Spirulina | Glycerine | None | control | 14 | 2914 | 3744 | 3306 |
| Spirulina | Glycerine | None | control | 15 | 2914 | 3744 | 3338 |
| Spirulina | Glycerine | Autoclaving | 1 cycle | 0 | 450 | 483 | 461 |
| Spirulina | Glycerine | Autoclaving | 1 cycle | 1 | 483 | 964 | 487 |
| Spirulina | Glycerine | Autoclaving | 1 cycle | 2 | 483 | 964 | 487 |
| Spirulina | Glycerine | Autoclaving | 1 cycle | 3 | 981 | 1011 | 994 |
| Spirulina | Glycerine | Autoclaving | 1 cycle | 4 | 1011 | 1090 | 1043 |
| Spirulina | Glycerine | Autoclaving | 1 cycle | 5 | 1091 | 1164 | 1113 |
| Spirulina | Glycerine | Autoclaving | 1 cycle | 6 | 1169 | 1262 | 1210 |
| Spirulina | Glycerine | Autoclaving | 1 cycle | 7 | 1262 | 1359 | 1336 |

| | | | | | | | |
|---|---|---|---|---|---|---|---|
| Spirulina | Glycerine | Autoclaving | 1 cycle | 8 | 1366 | 1450 | 1412 |
| Spirulina | Glycerine | Autoclaving | 1 cycle | 9 | 1498 | 1846 | 1638 |
| Spirulina | Glycerine | Autoclaving | 1 cycle | 10 | 1498 | 1846 | 1638 |
| Spirulina | Glycerine | Autoclaving | 1 cycle | 11 | 2021 | 2061 | 2051 |
| Spirulina | Glycerine | Autoclaving | 1 cycle | 12 | 2093 | 2134 | 2112 |
| Spirulina | Glycerine | Autoclaving | 1 cycle | 13 | 2174 | 2209 | 2188 |
| Spirulina | Glycerine | Autoclaving | 1 cycle | 14 | 2914 | 4000 | 3306 |
| Spirulina | Glycerine | Autoclaving | 1 cycle | 15 | 2914 | 4000 | 3338 |

**Table S2: mean pH of protein extracts n=3**

| Solvent | Process Parameter Changing to obtain protein | Parameter change | mean pH |
|---|---|---|---|
| Ultrapure water | Sonication | 12 cycles of 10s at 23kHz | 6.44 |
| Ultrapure water | Sonication | 18 cycles of 10s at 23kHz | 6.54 |
| Ultrapure water | Sonication | 24 cycles of 10s at 23kHz | 6.53 |
| Ultrapure water | Freeze-thawing | control | 6.63 |
| Ultrapure water | Freeze-thawing | 1 cycle 30 mins freeze, 45 mins thaw | 6.66 |
| Ultrapure water | Freeze-thawing | 2 cycle 30 mins freeze, 45 mins thaw | 6.64 |
| Ultrapure water | Autoclaving | control | 6.55 |
| Ultrapure water | Autoclaving | 1 cycle | 6.34 |
| PBS | Sonication | 12 cycles of 10s at 23kHz | 6.25 |
| PBS | Sonication | 18 cycles of 10s at 23kHz | 6.23 |

| | | | |
|---|---|---|---|
| PBS | Sonication | 24 cycles of 10s at 23kHz | 6.24 |
| PBS | Freeze-thawing | control | 6.47 |
| PBS | Freeze-thawing | 1 cycle 30 mins freeze, 45 mins thaw | 6.50 |
| PBS | Freeze-thawing | 2 cycle 30 mins freeze, 45 mins thaw | 6.49 |
| PBS | Autoclaving | control | 6.47 |
| PBS | Autoclaving | 1 cycle | 6.30 |
| 10% v/v Glycerin/ water solution | Sonication | 12 cycles of 10s at 23kHz | 6.57 |
| 10% v/v Glycerin/ water solution | Sonication | 18 cycles of 10s at 23kHz | 6.50 |
| 10% v/v Glycerin/ water solution | Sonication | 24 cycles of 10s at 23kHz | 6.52 |
| 10% v/v Glycerin/ water solution | Freeze-thawing | control | 6.63 |
| 10% v/v Glycerin/ water solution | Freeze-thawing | 1 cycle 30 mins freeze, 45 mins thaw | 6.66 |
| 10% v/v Glycerin/ water solution | Freeze-thawing | 2 cycle 30 mins freeze, 45 mins thaw | 6.66 |
| 10% v/v Glycerin/ water solution | Autoclaving | control | 6.55 |
| 10% v/v Glycerin/ water solution | Autoclaving | 1 cycle | 6.31 |
